# Formulation development and comparability studies with an aluminum-salt adjuvanted SARS-CoV-2 Spike ferritin nanoparticle vaccine antigen produced from two different cell lines

**DOI:** 10.1101/2023.04.03.535447

**Authors:** Ozan S. Kumru, Mrinmoy Sanyal, Natalia Friedland, John Hickey, Richa Joshi, Payton Weidenbacher, Jonathan Do, Ya-Chen Cheng, Peter S. Kim, Sangeeta B. Joshi, David B. Volkin

**Author notes:** Corresponding author: D.B. Volkin, Multidisciplinary Research Building, 2030 Becker Dr., Lawrence, KS 66047. Phone: -6262.

## Abstract

The development of safe and effective second-generation COVID-19 vaccines to improve affordability and storage stability requirements remains a high priority to expand global coverage. In this report, we describe formulation development and comparability studies with a self-assembled SARS-CoV-2 spike ferritin nanoparticle vaccine antigen (called DCFHP), when produced in two different cell lines and formulated with an aluminum-salt adjuvant (Alhydrogel, AH). Varying levels of phosphate buffer altered the extent and strength of antigen-adjuvant interactions, and these formulations were evaluated for their (1) *in vivo* performance in mice and (2) *in vitro* stability profiles. Unadjuvanted DCFHP produced minimal immune responses while AH-adjuvanted formulations elicited greatly enhanced pseudovirus neutralization titers independent of ∼100%, ∼40% or ∼10% of the DCFHP antigen adsorbed to AH. These formulations differed, however, in their *in vitro* stability properties as determined by biophysical studies and a competitive ELISA for measuring ACE2 receptor binding of AH-bound antigen. Interestingly, after one month of 4°C storage, small increases in antigenicity with concomitant decreases in the ability to desorb the antigen from the AH were observed. Finally, we performed a comparability assessment of DCFHP antigen produced in Expi293 and CHO cells, which displayed expected differences in their N-linked oligosaccharide profiles. Despite consisting of different DCFHP glycoforms, these two preparations were highly similar in their key quality attributes including molecular size, structural integrity, conformational stability, binding to ACE2 receptor and mouse immunogenicity profiles. Taken together, these studies support future preclinical and clinical development of an AH-adjuvanted DCFHP vaccine candidate produced in CHO cells.

## Introduction

The ongoing COVID-19 pandemic remains the largest worldwide infectious disease emergency since the Influenza pandemic of 1918-20. Despite the remarkable and rapid development of first-generation COVID-19 vaccines, therapeutics and nonpharmaceutical interventions, SARS-CoV-2 infections continue unpredictably as it mutates and circulates in both human and animal populations [1]. To date, Omicron subvariants continue to dominate globally and possess a definite reproductive advantage [2]. One major limitation of currently employed mRNA-based COVID-19 vaccines is their inability to generate sufficiently robust immune responses against SARS-CoV-2 variants. Furthermore, immunity induced both by vaccination and infection tends to wane over months as breakthrough infections and re-infections in the vaccinated and unvaccinated populations are common, respectively [3-5]. To address these gaps, development of second-generation COVID-19 vaccines with enhanced breadth and duration of protective immune responses is an ongoing high priority [6].

Another considerable limitation of currently available mRNA-based COVID-19 vaccines is their high costs, limited production capacity, and complicated supply chains including the requirement for ultralow temperature storage in freezers. Such attributes have resulted in limited global availability and poor vaccine coverage especially in low-and middle-income countries (LMICs). In particular, cold chain infrastructure for frozen storage is largely inadequate in many LMICs, which makes widespread distribution of mRNA vaccines challenging. Therefore, there is also an urgent need to supply the world with low-cost, second generation COVID-19 vaccines that can be easily manufactured, distributed, and stored in the refrigerator (or even under ambient conditions) [7, 8].

Various alternative vaccine platforms for developing next-generation COVID-19 vaccine candidates with improved efficacy that will also enable improved vaccine coverage in LMICs are currently being explored [9]. Recombinant protein subunit vaccines are an attractive approach due to the relative ease and scalability of antigen production combined with comparable or superior immunogenicity profiles compared to viral vectored and mRNA vaccines [10]. One such approach is expression of the SARS-CoV-2 spike protein fused to *H. pylori* ferritin (non-heme) that subsequently self-assembles as a nanoparticle that enables multivalent antigenic display. Other vaccine candidates have utilized this approach with different antigens including influenza HA, HIV env, and *Borrelia burgdoferi* OspA [11-13]. Previous and ongoing work demonstrate the prefusion SARS-CoV-2 spike protein fused to *H. pylori* ferritin induce high levels of neutralizing antibodies in mice and non-human primates [14-16]. In this work, we examined an updated version of the spike-ferritin-nanoparticle construct termed Delta-C70-Ferritin-HexaPro or DCFHP, to which four additional stabilizing proline mutations were introduced to the previously described 2P construct both of which include a 70 amino acids C-terminal truncation [14, 17]. This updated version of the nanoparticle-based vaccine candidate, when formulated with aluminum hydroxide as the sole adjuvant, elicited potent and durable neutralizing antisera in mice and NHPs SARS-CoV-2 Wuhan-1 and several variants of concern, as well as against SARS-CoV-1 [17].

Protein subunit antigens generally require adjuvants to enhance immune responses [18, 19], and Powell et al. (2021) [14] previously reported that the 2P construct of spike-ferritin-ΔC nanoparticle construct (SΔC-Fer), when formulated with Quil-A and MPLA as adjuvants (prime+boost), generated high levels of neutralizing antibody titers in mice [14]. The ability of DCFHP antigen formulated with aluminum-salt adjuvants, with and without CpG oligonucleotides, to generate similarly robust immune response in mice and NHPs was recently reported by [17]. Formulations adjuvanted solely with aluminum-salts (alum), however, are ideal for use in LMICs since they are inexpensive, easy to produce at large-scale, and their clinical use is supported by decades of safety and efficacy data in childhood, adolescent and adult vaccines [20].

The focus of this work was formulation development and analytical characterization of DCFHP with aluminum salt adjuvants to facilitate future preclinical and clinical development as a COVID-19 vaccine candidate targeted for use in LMICs. We evaluated and optimized DCFHP formulations with aluminum-salt (alum) adjuvants both in terms of *in vivo* performance and *in vitro* pharmaceutical properties. First, we determined the effect of varying the extent and strength of antigen-adjuvant interactions on mouse immunogenicity and storage stability. To this end, we developed a relative binding assay (competitive ELISA format) to measure binding of AH- adsorbed DCFHP antigen to the ACE2-receptor. Next, we evaluated the DCFHP antigen with a variety of physiochemical methods and identified sensitive analytical tools to define potential critical structural attributes (i.e., primary structure, post-translational modifications, higher-order structural integrity, and molecular size including aggregation). Finally, we performed a comparability assessment using a combination of physicochemical, ACE2 receptor binding, and mouse immunogenicity studies to compare DCFHP produced in Expi293 and CHO cells. The practical implications of these results to guide future CMC (chemistry, manufacturing, control) development of a low-cost, aluminum-salt adjuvanted DCFHP as a subunit COVID-19 vaccine candidate are discussed.

## Materials and Methods

### Materials

Spike-Ferritin nanoparticles (DCFHP) were previously described by Powell et al., [14], with exception of four additional proline amino acid substitutions F817P, A892P, A899P, A942P as described in [21]. The DCFHP nanoparticles were expressed in Expi293 cells as described previously [14], with minor modifications as described in the Supplemental methods. CHO cell-based production of DCFHP is also described in the Supplemental methods.

### Methods

The analytical methods employed in this work including physicochemical techniques (i.e., circular dichroism, intrinsic fluorescence spectroscopy, differential scanning calorimetry, SEC-MALS, sedimentation velocity-AUC, dynamic light scattering, LC-MS peptide mapping, N-linked oligosaccharide mapping) as well as immunochemical binding and *in vivo* potency assays (i.e., bio-layer interferometry, competitive ELISAs, and mouse immunogenicity studies including pseudovirus neutralization titers) are described in detail in the Supplemental methods. In addition, various experimental setups used for antigen-adjuvant binding (e.g., Langmuir isotherms), antigen-adjuvant desorption, and vaccine stability studies are each provided in the Supplemental methods.

## Results

### DCFHP antigen-Aluminum-salt adjuvant interaction studies

First, we determined the extent of DCFHP antigen binding to aluminum hydroxide (Alhydrogel™, AH) and aluminum phosphate (Adjuphos™, AP) adjuvants. Both AH and AP are colloidal suspensions that can adsorb protein antigens by a variety of molecular mechanisms including for electrostatic, hydrophobic and covalent interactions [22]. We observed antigen 100% binding to AH (**Figure 1A-B**) and incomplete binding to AP (data not shown). These results are consistent with electrostatic interactions between the negatively charged DCFHP (calculated pI ∼5.9) and the positively charged AH particles (zeta potential ∼ +30 mV) at neutral pH [23]. The partial binding of DCFHP to AP particles (zeta potential ∼ −30 mV at neutral pH), however, is likely driven by non-electrostatic forces such as hydrophobic interactions [24].

**Figure 1.**
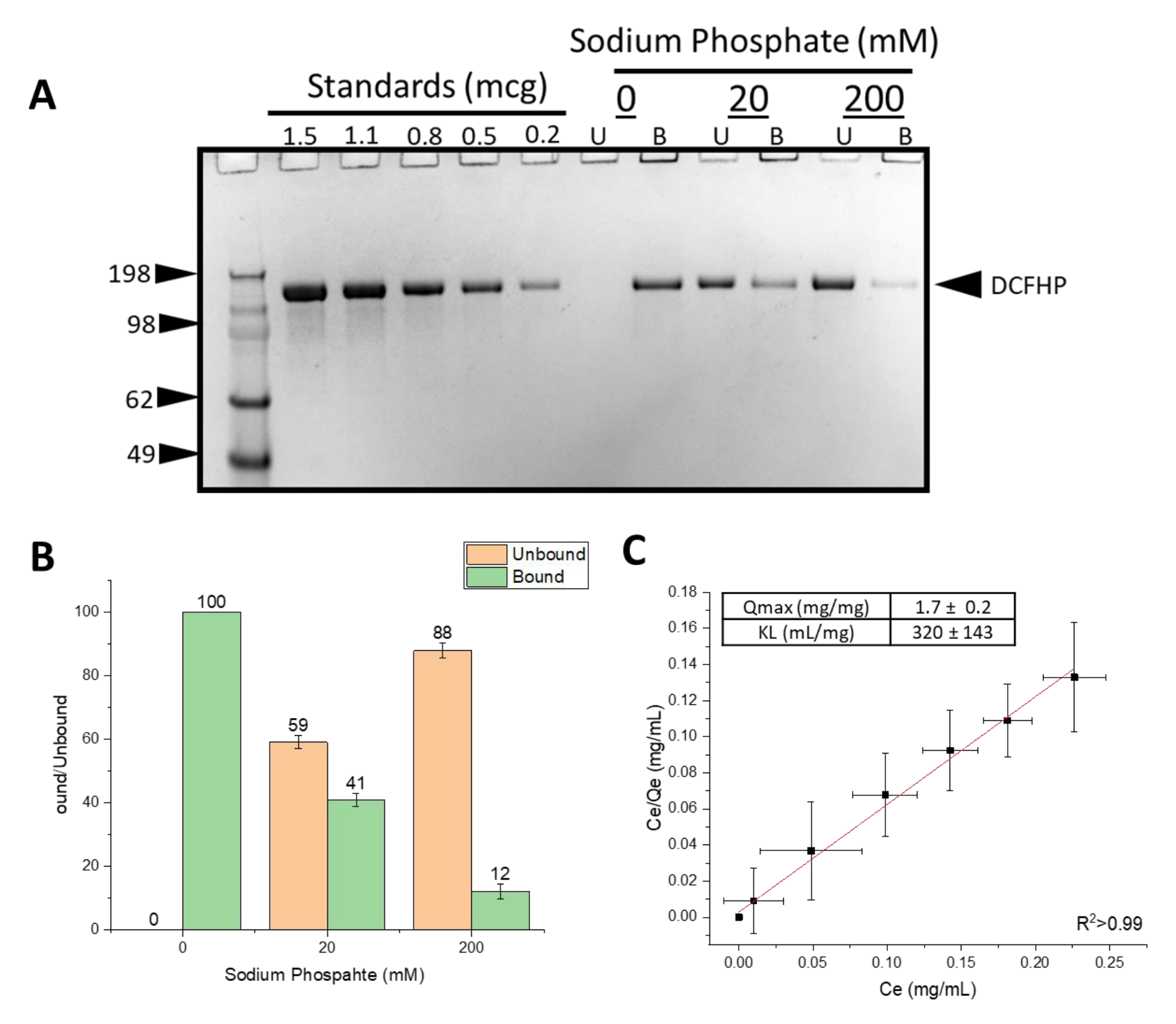
Binding of DCFHP antigen to Alhydrogel adjuvant in the presence and absence of varying levels of phosphate buffer. (A) representative SDS-PAGE gel where the AH-bound fractions were determined by strong desorption of antigen from the AH adjuvant (see text). The effect of 0, 20mM and 200mM sodium phosphate buffer on the binding of DCFHP to Alhydrogel (AH) was determined by centrifugation of samples followed by SDS-PAGE analysis and estimation of the amount (b) bound and (U) unbound in each fraction by densitometry. (B) Percent of DCFHP antigen bound to AH with indicated phosphate buffer concentrations in formulations containing 50 mcg/mL DCFHP, 50 mcg AH in 20mM Tris, 150mM NaCl, 5% Sucrose, pH 7.5. Values are the mean of five independent experiments with error bars representing the standard deviation. (C) Langmuir isotherm analysis antigen binding (DCFHP with no sodium phosphate) to AH with the linear fit of adsorption data displayed in red with adsorption capacity and strength values displayed in box (see methods section for details). Langmuir isotherm data were the mean of four separate adsorption isotherms and the error bars represent the standard deviation.

Pre-treating AH with varying concentrations of sodium phosphate alters the overall surface charge of the adjuvant from positive to negative by ligand exchange of a phosphate for hydroxyl on the surface of the aluminum salt [24, 25]. Pre-treatment of AH with 0, 20 and 200 mM sodium phosphate changed the AH surface charge (zeta potential) from positive to negative, which in turn decreased the amount of DCFHP bound to AH from ∼100%, ∼40% and ∼10%, respectively (**Figure 1A-B**). Increasing the sodium phosphate concentration even further (>200mM), however, did not further affect the desorption of DCFHP from AH (data not shown), a result also consistent with some non-electrostatic interactions between both alum adjuvants and DCFHP.

To better define the extent and strength of AH-DCFHP interactions, we performed adsorption isotherm experiments in the absence of phosphate buffer (see Supplemental methods). The data fit well to the Langmuir equation showing monolayer adsorption at ∼1.7 mg DCFHP/mg AH (**Figure 1C**), which is about an order of magnitude higher binding capacity than the AH- adsorbed DCFHP doses used in murine studies (see below). The adsorptive strength (K_L_) value of the antigen-adjuvant interaction was measured as ∼320 mL/mg), a value on the high-end of the range observed with other recombinant protein antigens with AH [23] (see Discussion).

### Effect of adjuvant-antigen interactions on mouse immunogenicity profiles of Alhydrogel-adjuvanted DCFHP

The effect of varying the extent of DCFHP binding to AH on SARS-CoV-2 neutralizing immune responses in mice was determined using an established prime/boost model [14], with four different groups of mice immunized with 10 µg DCFHP in each formulation (**Figure 2**). First, formulations that did not contain AH adjuvant generated ∼2 log lower pseudoviral neutralization titers at days 42 and 63, which confirms an adjuvant is required to generate a robust immune response (Figure 2). For AH containing formulations with and without pretreatment with 20 mM and 200 mM sodium phosphate, no differences in pseudovirus neutralization titers (Wuhan-1) were observed at day 21 (pre-boost), and at days 42 and 63 (+booster dose at day 21) (Figure 2). No changes in the neutralization titers were observed at day 63, which indicates durability of the neutralizing immune response (Figure 2). Taken together, these results show that generation of a sufficient neutralizing antibody response requires the presence of AH adjuvant but is independent of the amount of DCFHP bound to AH adjuvant and independent of sodium phosphate pre-treatment to modulate the binding extent and strength.

**Figure 2.**
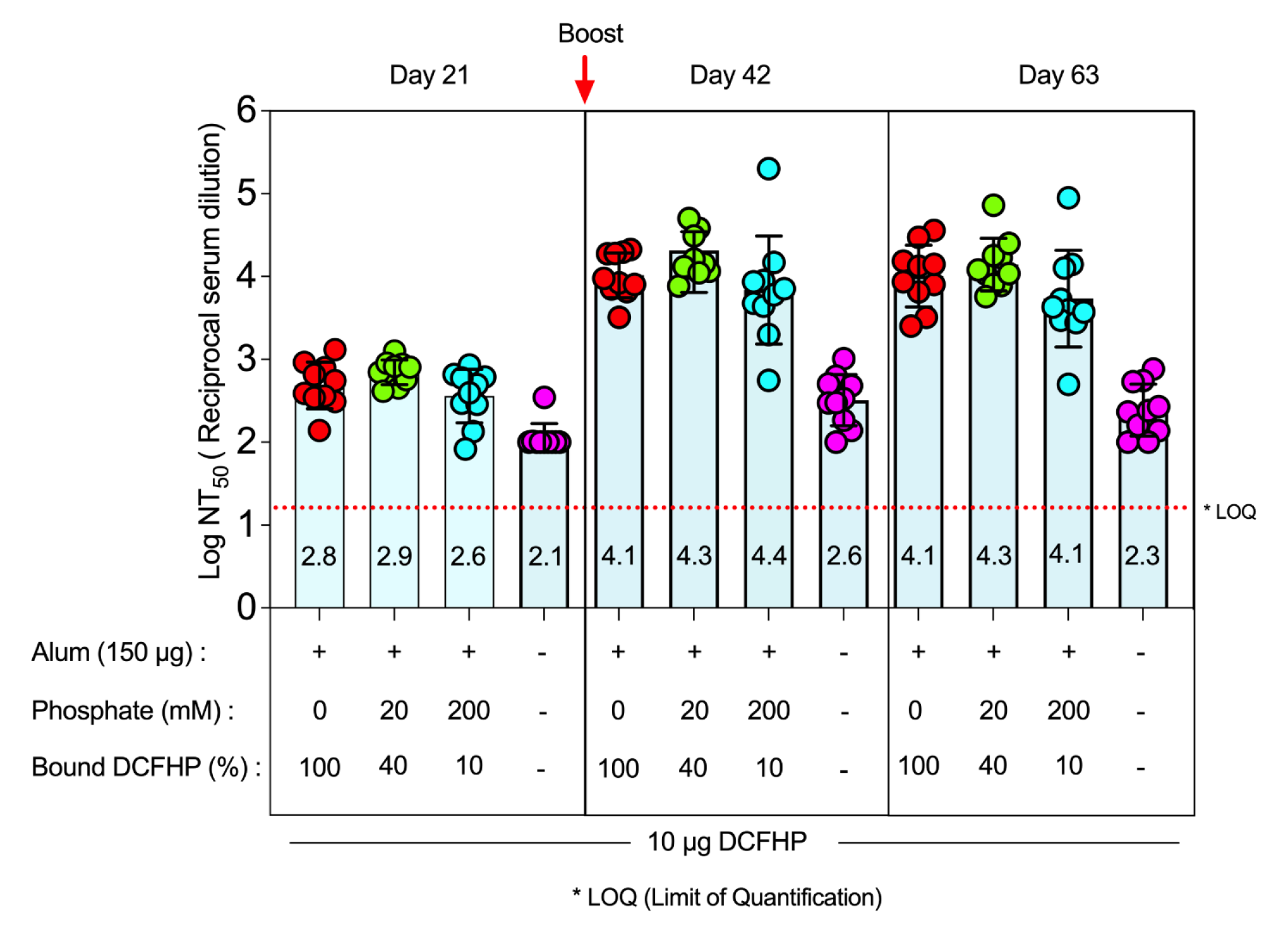
Pseudovirus neutralization titers in mice after immunization with various formulations of DCFHP. Mice were immunized (prime dose at day 0, booster dose at day 21) with 10 mcg of DCFHP antigen, 150 mcg Alhydrogel (alum) adjuvant in a formulation buffer (20mM Tris, 150mM NaCl, 5% Sucrose, pH 7.5) with the indicated sodium phosphate concentrations. The percent of DCFHP antigen bound to alum adjuvant is also indicated. The injection volume was 0.1 mL. Individual titers of ten mice are displayed and the bars represent the mean, and the error bars represent the 95% confidence interval. * LOQ, Lower limit of Quantitation.

### Effect of adjuvant-antigen interactions on in vitro structural integrity and stability of AH-adjuvanted DCFHP

Next, to evaluate how the interactions between the DCFHP antigen and AH-adjuvant affects the pharmaceutical properties of the vaccine candidate, the formulated AH-DCFHP samples were evaluated (and compared to antigen in solution without AH) for conformational stability (differential scanning calorimetry, DSC), antigen binding to ACE2 receptor (competitive ELISA), and the ability to desorb the antigen from AH (mild vs strong desorption conditions; see Supplemental methods), with the latter two evaluated both at time zero and after one month of storage at 4° and 25°C.

First, DSC analysis of the DCFHP antigen in solution displayed multiple thermal unfolding events (Tm1, Tm2, Tm3) at ∼50, ∼63, and ∼83°C (**Figure 3A**), respectively, a result consistent with multidomain nature of the DCFHP antigen. Upon antigen adsorption to AH that was pre-treated with 0, 20 or 200 mM sodium phosphate (i.e., 100%, 40% to 10% DCFHP bound to AH, respectively), the various AH-adsorbed DCFHP antigen showed a decrease in only the Tm3 value (from ∼83 to ∼74°C) (**Figure 3B-E**). Interestingly, there was a concomitant notable reduction in the apparent enthalpy (ΔH’) values for AH-bound antigen samples (**Figure 3F**). Taken together, these DSC results are consistent with DCFHP-AH interactions altering the overall conformational stability of the antigen as observed by changes in Tm3 and ΔH’ values. When the AH was pre-treated with 20- or 200-mM sodium phosphate, interactions between AH and DCFHP are modified resulting in ΔH’ values between those observed for the antigen in solution and completely AH-adsorbed (with Tm3 values remaining decreased).

**Figure 3.**
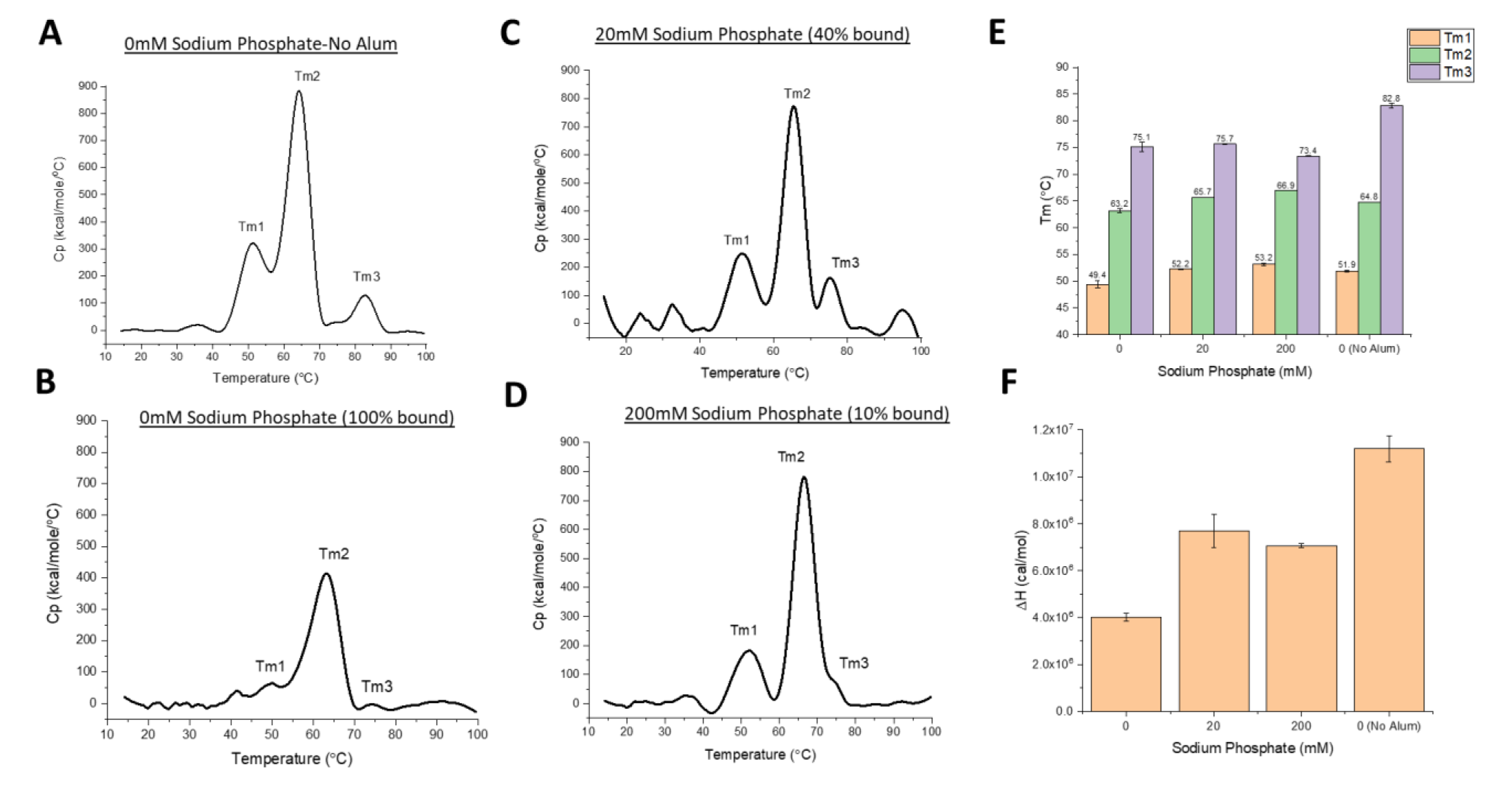
Differential scanning calorimetry analysis of DCFHP formulations in the presence and absence of Alhydrogel adjuvant with varying concentrations of sodium phosphate buffer. Representative thermograms of DCFHP (A) in solution no sodium phosphate and no Alhydrogel (AH), (B) no sodium phosphate and AH-adsorbed, (C) 20 mM sodium phosphate and AH-adsorbed, and (D) 200mM sodium phosphate and AH-adsorbed. The percent antigen bound to AH is shown in header of each panel. Analysis of DSC data from each sample included determination of (E) thermal melting temperature (Tm) values, and (F) apparent enthalpy (ΔH’) values. DCFHP samples were formulated in a buffer containing 20mM Tris, 150mM NaCl, 5% Sucrose, pH 7.5 with the indicated sodium phosphate concentrations. Representative thermograms are displayed and data are average of two duplicate scans with the error bars representing the data range.

Second, we determined how antigen-adjuvant interactions affect the local structural integrity of the key epitopes in the antigen by employing *in vitro* binding assays that monitor DCFHP binding to the ACE2 receptor. Since BLI measurements are typically unsuitable for direct analysis of AH-bound antigens (i.e., requires antigen desorption or dissolving the adjuvant), we developed a competition ELISA assay to assess the ability of ACE2 receptor to bind DCFHP when AH-adsorbed. This assay format monitors the binding the ACE2 to the DCFHP antigen when formulated in solution or AH-absorbed with no notable differences (**Figure 4A**). To establish the assay is stability-indicating, we incubated AH-adsorbed DCFHP samples overnight at 4°, 40°, and 50°C and measured the ACE2 binding (**Figure 4B-C**). While a small increase in ACE2 binding was observed at 4°C (see below), a notable loss (∼60% and ∼80%) was observed at elevated temperatures (40 and 50°C, respectively).

**Figure 4.**
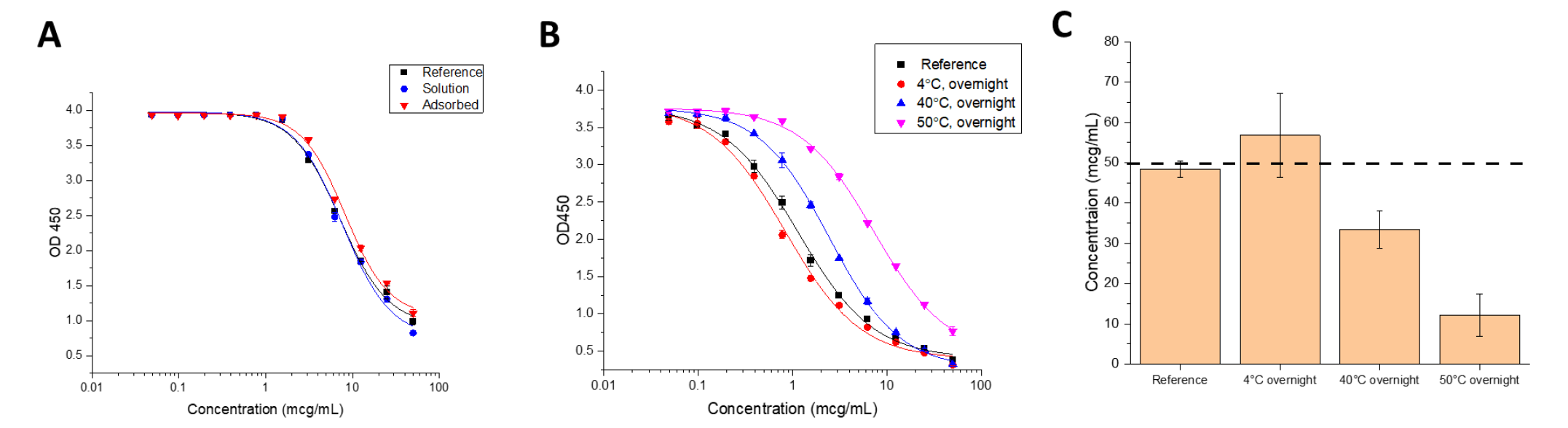
Competitive ELISA measuring binding of ACE2 receptor to DCFHP samples in solution and adsorbed to Alhydrogel adjuvant. (A) Representative ELISA curves comparing ACE2 binding of DCFHP while in solution and adsorbed to Alhydrogel. (B) Representative ELISA curves of AH-adsorbed DCFHP incubated at 4°, 40°, and 50°C overnight and compared to a DCFHP reference sample that was adsorbed to Alhydrogel on the same day of the assay. (C) Native antigen concentration of thermally-stressed AH-adsorbed DCFHP samples as determined from the ELISA curves (panel B) versus the reference sample. Data are the mean of two independent experiments, which contained two replicates (n=4) with the error bars representing the standard deviation. The dashed line represents the target concentration of 50 mcg/mL. DCFHP samples were at 50 mcg/mL in 20 mM Tris, 150mM NaCl, 5% Sucrose, pH 7.5 buffer, with and without being adsorbed to 1.5 mg/mL Alhydrogel.

We then employed the competitive ELISA to evaluate DCFHP samples used in the mouse immunogenicity study, including injections at time zero (day prior to mouse immunization) and after 1 month of incubation at 4°C (booster dose administered after 21 days), We also assessed thermally stressed samples (1 month at 25°C). Interestingly, while all samples displayed similar ACE2 binding at time zero (**Figure 5A**), the solution and AH-adsorbed samples showed a net increase in the levels of ACE2 binding after storage for 1 month at 4°C (while the two phosphate pretreated AH samples showed no change during storage). After storage at 25°C for one month, however, we observed a consistent decrease *in vitro* ACE2 binding with the largest decrease observed in DCFHP samples in solution and formulated with AH + 200 mM sodium phosphate (Figure 5A).

**Figure 5.**
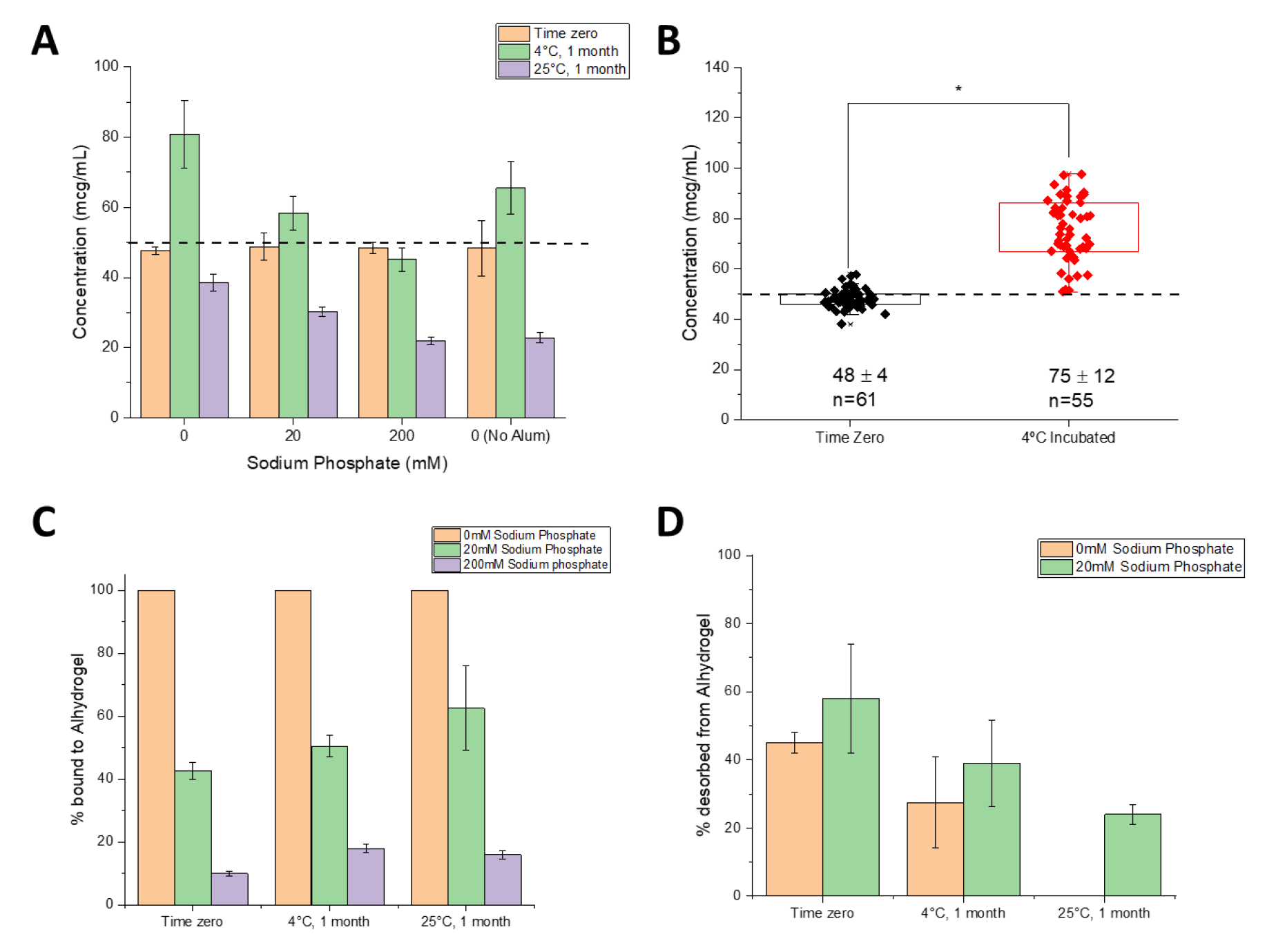
Nature of the interaction of DCFHP antigen with Alhydrogel adjuvant as a function of storage time, temperature, and sodium phosphate concentration. (A) ACE2 competitive ELISA results for the DCFHP formulations (three AH-adsorbed and one without alum as indicated) used for mouse immunization studies at time zero (first dose) and after 1 month storage at 4 (second dose) as well as additional samples stored at 25°C for 1 month. Data are the mean of two independent samples, each measured twice with the error bars represent the standard deviation. (B) Box plot analysis of ACE2 competitive ELISA data from 21 independent experiments of DCFHP antigen 100% adsorbed to Alhydrogel (no phosphate buffer) at time zero and after incubation at 4°C for an average of 10 days (range 3-21 days). *Results from a two tailed student’s t-test indicate the two data sets are significantly different (*p<0.0001). The dashed line represents the target concentration of 50 mcg/mL DCFHP. (C, D) Binding strength of DCFHP to AH increases as a function of storage time and temperature. (C) The percent of DCFHP bound to Alhydrogel at time zero and after 1 month incubation at 4 and 25°C as determined by strong desorption (0.4 M sodium phosphate, LDS, 95°C incubation for 5-10 min), and (D) the percent of DCFHP desorbed from same samples using mild desorption conditions (0.4 M sodium phosphate, 37°C incubation for 30 min). Data are the mean of two replicates with the error bars representing the data range.

To further explore the observation of increased ACE2 receptor binding during storage at 4°C, we analyzed competitive ELISA data (from an aggregate of >20 independent experiments) by comparing AH-adsorbed DCFHP samples that were stored at 4°C for 3-21 days (mean incubation time of 10 days) versus DCFHP samples that were freshly adsorbed to AH after thawing from −80°C (time zero). Over 50 individual data points were analyzed statistically using a box-whisker analysis (**Figure 5B**), and two notable trends were observed: (1) there was more variability in the samples stored at 4°C (75 ± 12 mcg/mL) when compared to the freshly adsorbed samples at time zero (48 ± 4 mcg/mL), and (2) there was a statistically significant ∼1.6X increase in the ability to bind ACE2 receptor after storage at 4°C (p<0.0001 by two-tailed students t-test); see Discussion.

Finally, it has been previously reported that AH-adsorbed subunit antigens stored over time display a decreasing ability to desorb the antigen from the adjuvant [22, 25]. In this work, we employed two different desorption procedures to explore the effect of storage time and temperature on interactions between DCFHP antigen and AH adjuvant (see Supplemental methods): (1) a “strong desorption” procedure under denaturing conditions (combination of phosphate buffer, LDS, DTT and heat), and (2) a “mild desorption” procedure under non-denaturing conditions (phosphate buffer only without detergent, reducing agent or heat). Using strong desorption conditions, the percent DCFHP bound to AH-surface was determined at time zero (∼100, ∼40, and ∼10% antigen bound to AH at 0, 20, and 200 mM phosphate, respectively), and these percent bound values did not change after storage for 1 month at 4 and 25°C (**Figure 5C**). This result demonstrates, that across all AH-adsorbed DCFHP samples, ∼100% of antigen was desorbed from AH after storage using strong desorption conditions. In contrast, using mild desorption conditions with 100% AH-bound antigen sample (no phosphate buffer) stored over one month, only ∼50% of DCFHP was desorbed from AH at time zero, while the percent of DCFHP desorbed from AH decreased to ∼25% and ∼0% after one month at 4° and 25°C, respectively (**Figure 5D**). Similar results of decreasing ability to desorb antigen from AH adjuvant by mild desorption during storage was observed in the AH-DCFHP formulation containing 20mM sodium phosphate (Figure 5D), albeit less antigen was AH-bound at time zero (similar experiments with the 200 mM phosphate samples were not performed since ∼90% of the antigen was desorbed from AH at time zero).

### Comparability assessment of Expi293 and CHO produced DCFHP nanoparticles

The formulation and characterization work described above was performed with small amounts of DCFHP antigen produced at the lab scale using Expi293 cells. Before additional formulation and longer-term stability studies could be considered for future work (see Discussion), we assessed the quality of the DCFHP antigen produced in Expi293 and CHO cells, the latter to be used to enable larger scale production. To this end, we employed various physiochemical methods to measure primary structure, post-translational modifications, molecular size, aggregation, and higher-order structural integrity of the protein antigen as well as *in vitro*-derived ACE2 binding studies and *in vivo* immunogenicity in mice (pseudovirus neutralization titers). The primary structure of the DCFHP antigen produced in the two cell lines was assessed by LC-MS peptide mapping (**Figure 6A, B**). The relative base-peak ion abundance chromatograms of PNGaseF-treated and trypsin or chymotrypsin digested CHO and Expi293 DCFHP samples were overall similar with no new or missing peaks (some minor retention time shifts (< 2 min) were observed between the two samples, likely due to the method’s long elution gradient (>70 min). Following peptide identification using MS/MS, the overall sequence coverage of CHO or Expi293 DCFHP was 94% and 92%, respectively. The sequence coverage of the Spike portion and Ferritin portion of each polypeptide was 93% and 98% for CHO DCFHP, and 91% and 98% for Expi293 DCFHP (**Supplemental Figure S1**). Although additional method development is required for complete sequence coverage, the fingerprint analysis of the peptide mapping chromatograms demonstrates the primary sequence of DCFHP protein produced from the two cell lines are overall similar with no differences in potential chemical degradation byproducts such as Met oxidation.

**Figure 6.**
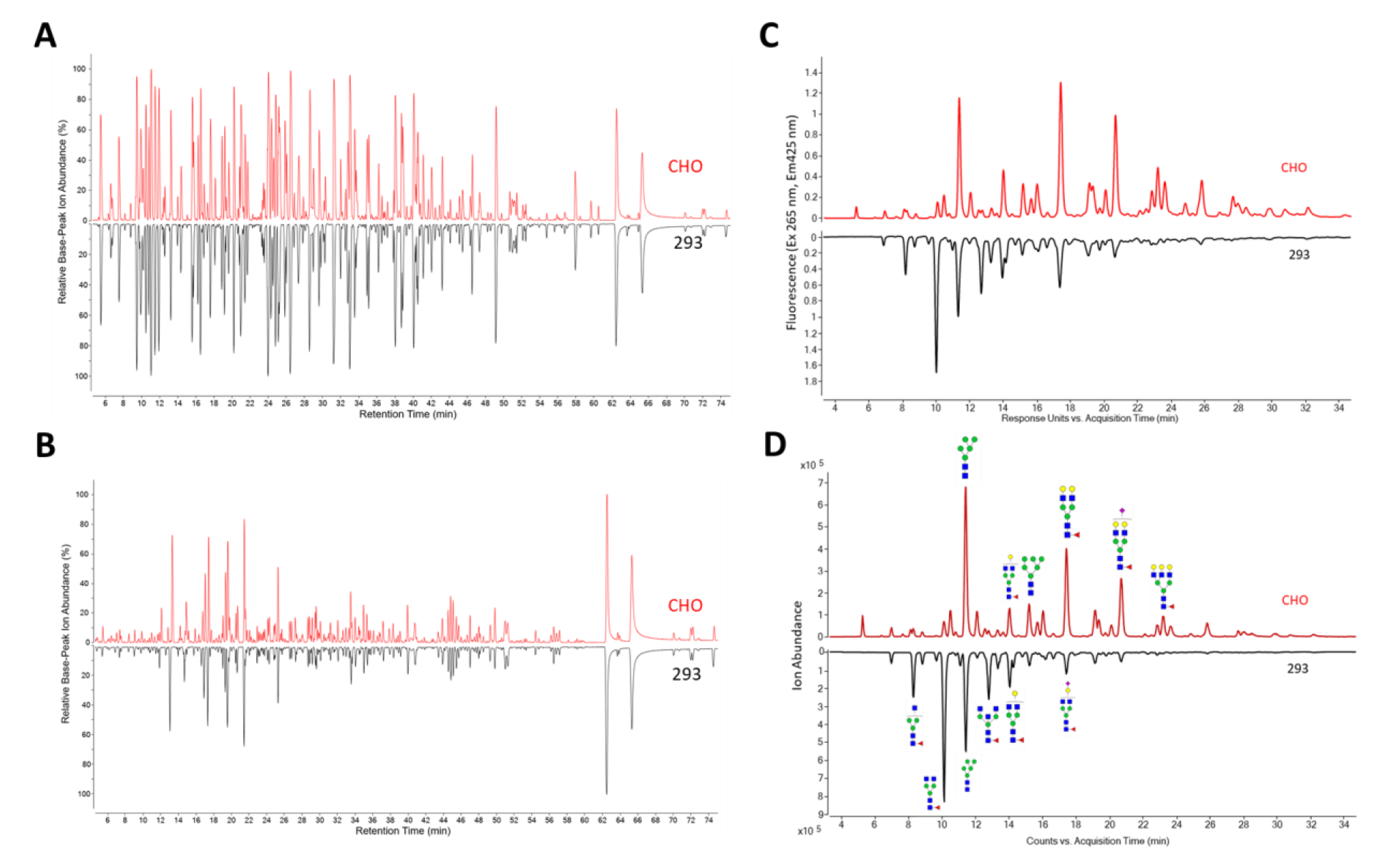
Comparison of primary structure and post-translational modification of DCFHP produced in Expi293 and CHO cells as measured by LC MS peptide mapping and N-linked oligosaccharide mapping. Representative LC-MS peptide mapping chromatograms of DCFHP (pre-treated with PNGaseF) and digested with either A) Trypsin or B) Chymotrypsin. (C, D) Representative oligosaccharide mapping results of labelled N-glycans (removed by PNGaseF treatment) from DCFHP produced in Expi293 and CHO cells. C) fluorescence detection chromatograms or D) total ion count MS detection chromatograms. Data is representative from triplicate experiments.

N-linked glycan profiles of the Expi293 and CHO produced proteins were determined by oligosaccharide mapping and displayed notable differences. The CHO DCFHP consisted of more complex and higher sialic acid content while the Expi293-derived DCFHP contained more higher-ordered mannose N-glycans (**Figure 6C-D**, **Supplemental Tables S1, S2**). The fluorescence (Figure 6C) and total ion counts (Figure 6D) indicated the most abundant N-linked glycan in the 293 and CHO produced DCFHP were H3N4F1 and H5N4F1, respectively (Figure 6C-D, Supplemental Tables S1-S2). These results are consistent with observations made by other groups with regards to the spike trimer [26].

Molecular size determination of DCFHP was assessed by SEC-MALS, sedimentation SV-AUC and DLS. No notable differences were observed in terms of molecular size between the Expi293 and CHO produced nanoparticles. SEC-MALS results (**Figure 7A**) were consistent with previous observations [14] with the main nanoparticle species eluting between 8-10 minutes and a minor species eluting at 6-8 min, which presumably represents soluble higher molecular weight aggregates (Figure 7A). SV-AUC analysis revealed one main species (∼40S) and two minor species likely representing incompletely formed (∼25s) and agglomerated nanoparticles (**Figure 7B**). The smaller species likely eluted as part of the nanoparticle species on SEC-MALS and was only detected by SV-AUC. Similarly, DLS data (by intensity distribution) confirmed a fully formed nanoparticle of ∼40 nm and did not display notable aggregates in the range of 100-1000 nm (**Figure 7C**) for both Expi293 and CHO produced material.

**Figure 7.**
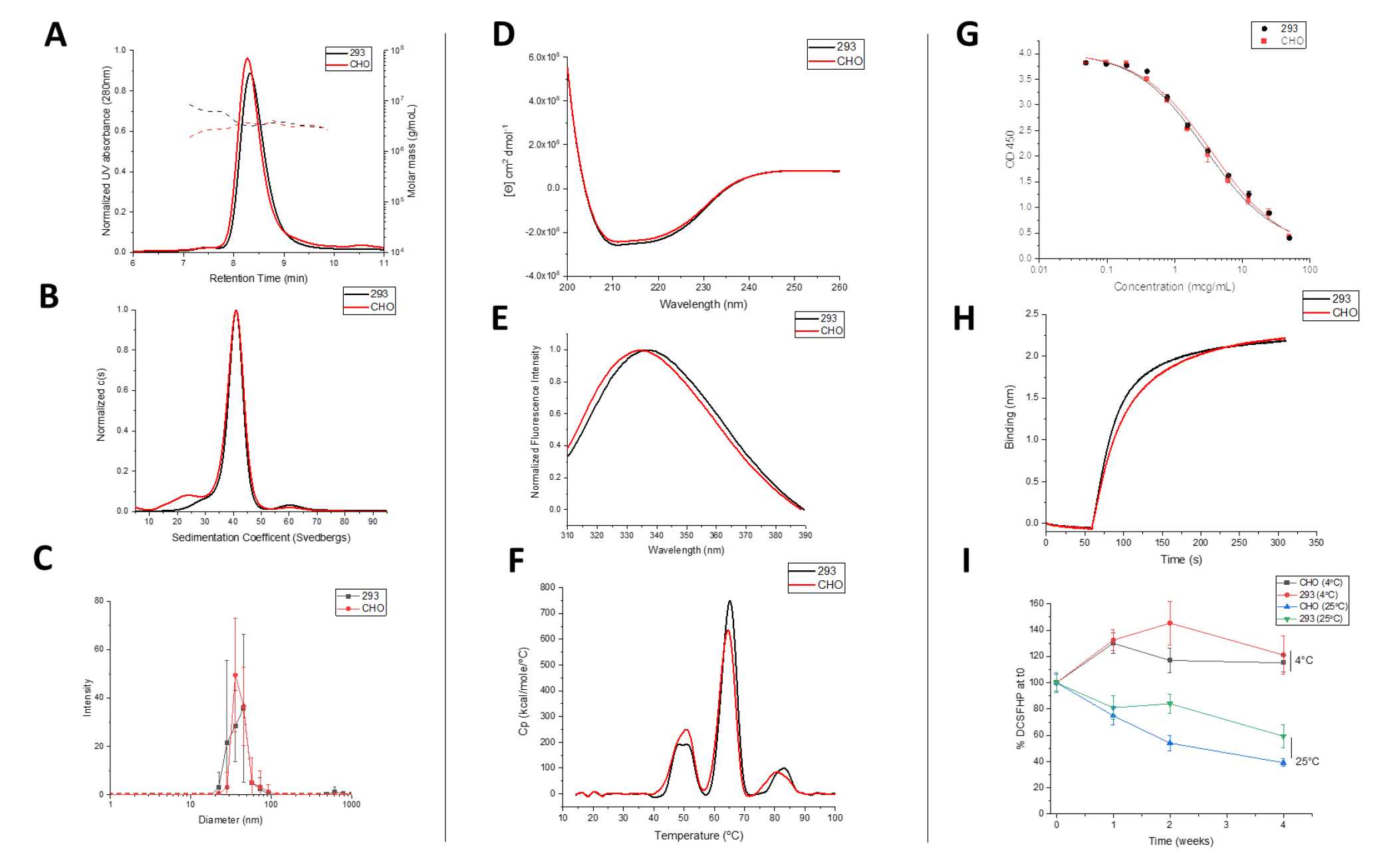
Physiochemical characterization of DCFHP produced in Expi293 and CHO cells as measured by biophysical and ACE2 receptor binding studies. Molecular size characterization in (A) SEC-MALS, (B) SV-AUC and (C) DLS. Overall secondary and tertiary structural integrity analysis at 10°C as measured by (D) circular dichroism, and (E) intrinsic fluorescence spectroscopy, respectively, and overall conformational stability profile as determined by (F) DSC. Representative binding curves of each DCFHP antigen to ACE2 as measured by (G) competitive ELISA, and (H) BLI. (I) Relative ACE2 binding (normalized to time zero) as measured by competitive ELISA in 293 and CHO produced DCFHP samples adsorbed to AH as a function of storage temperature (4 and 25°C) and time (up to 4 weeks). Samples were formulated in 20mM Tris, 150mM NaCl, 5% Sucrose, pH 7.5 buffer. Data are the mean of at least three independent measurements with the error bars representing the standard deviation. Representative data are shown in panels A, D, G, and H with the mean data are displayed in panels B, C, E, F, G and I.

CD analysis of both DCFHP samples revealed predominantly α-helical content from the double minima observed at ∼208 and ∼222 nm (**Figure 7D**). Similarly, the overall tertiary structure analysis of both DCFHP samples by intrinsic tryptophan fluorescence spectroscopy revealed a similar peak emission maximum of 336-337 nm, which indicates the average tryptophan residue is relatively solvent inaccessible (**Figure 7E**). The DSC profiles of the two samples showed similar three thermal transitions with Tm values of ∼49, ∼64, and ∼83°C (**Figure 7F**). No thermal unfolding events were observed when ferritin nanoparticles not fused to the spike trimer were tested (data not shown) indicating these signals are related to structural alterations in the spike protein. Compared to previously reported results for the thermal unfolding behavior of the spike trimer alone, where two unfolding events were observed at 49 and 64°C [27], the DSC results with DCFHP differ in terms of the additional third unfolding event was relatively weak and potentially is due to the spike protein incorporated into a nanoparticle or differences in solution conditions. In summary, no differences in higher-order structural integrity and conformational stability of the DCFHP produced from Expi293 and CHO produced cells.

In terms of *in vitro* binding assay, no notable differences in the ability of the two DCFHP samples to bind ACE2 was determined by competitive ELISA and BLI (**Figure 7G-H**, respectively). Negligible dissociation was observed in the BLI experiments (data not shown), a result which indicates strong binding of DCFHP to ACE2 receptor, as has been previously observed with another SARS-CoV-2 RBD nanoparticle vaccine candidate [28]. We also performed a 4-week stability study to measure the relative ACE2 binding ability of the CHO and 293 produced DCFHP antigens adsorbed to AH adjuvant (4 and 25°C) (**Figure 7I**). No notable differences in ACE2 binding ability were observed at 4°C. Similarly, no statistical differences were observed after 4 weeks at 25°C between the two samples (i.e., the 95% CI bands of the linear regression slopes overlapped; data not shown). Taken together, no notable differences in ACE2 receptor binding and relative storage stability profiles were observed between the DCFHP antigen produced in CHO and Expi293 cells.

Finally, no differences in *in vivo* immunogenicity were observed as measured by SARS-CoV-2 pseudovirus neutralization titers in mice immunized with either Expi293 or CHO produced DCFHP adsorbed to AH adjuvant (**Figure 8A-B**). Approximately ∼1 log higher titers were observed in neutralization of Wuhan-1 strain after a booster dose (Figure 8A). Conversely, neutralization titers of the Omicron variant (B.1.1.529.1) were ∼1.5-2 log lower when compared to Wuhan-1 strain (Figure 8B). The reduced neutralizing activity of the Omicron variant has been observed with other subunit vaccines after immunization with the Wuhan-1 spike protein [29].

**Figure 8.**
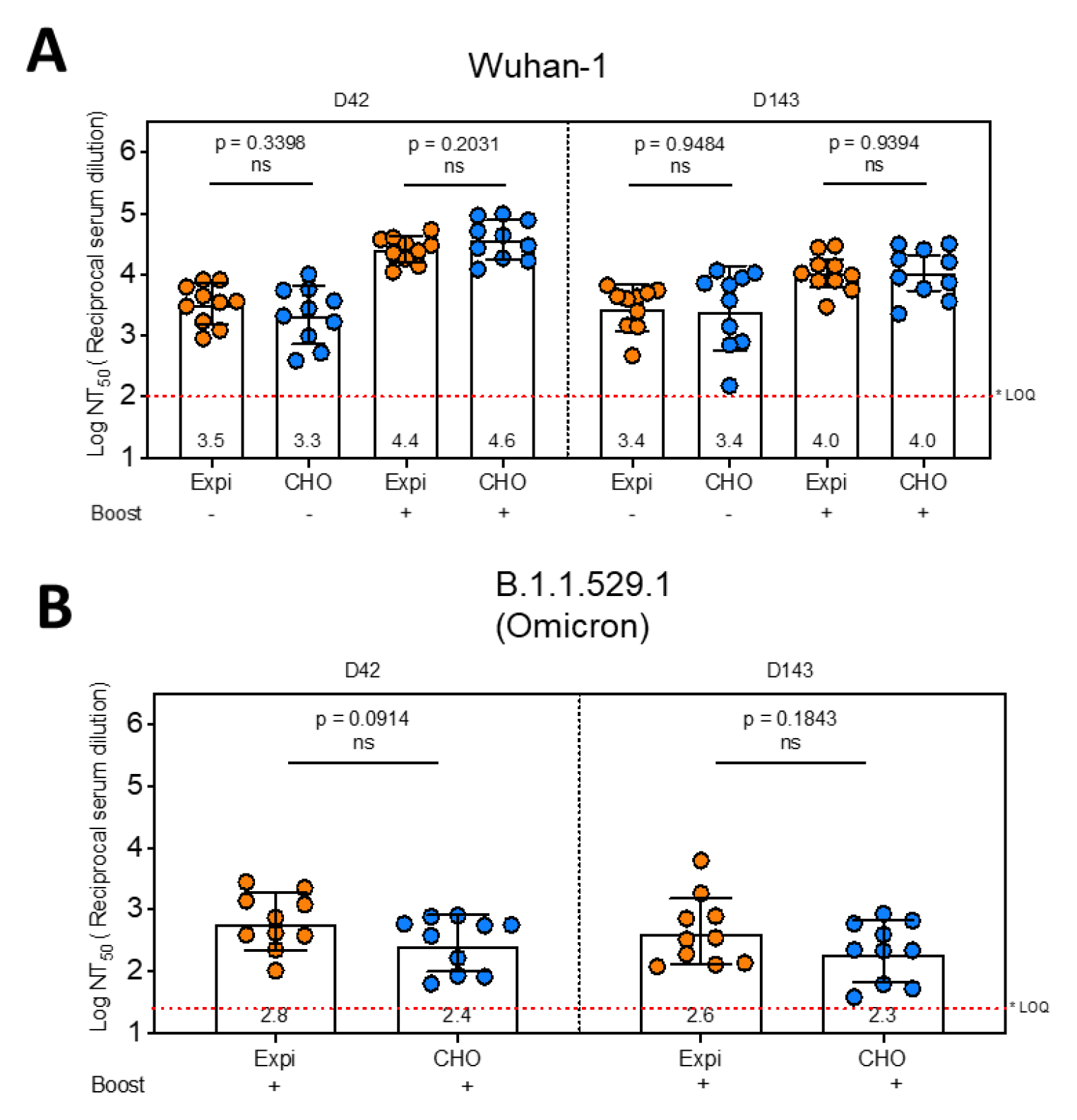
Pseudovirus neutralization titers of AH-adsorbed DCFHP produced in Expi293 and CHO cells. Mice were immunized with 10 mcg DCFHP, produced in either Expi-293 (transient expression) or CHO (stable expression) along with 150 mcg of alum (Alhydrogel). Some mice were boosted 21 days after the primary immunization as indicated in the figure. Blood samples were collected at indicated time points and the neutralization capacity of the serum samples were analyzed for SARS-CoV-2 wild type (A) and B.1.1.529.1 (Omicron) variant (B) using pseudovirus and HeLa cells co-expressing ACE2 and TMPRSS2. The neutralization titers are expressed as log_10_ values of 50 percent of neutralization titers (NT_50_). Note that the neutralization capacity is indistinguishable among the groups immunized with Expi-DCFHP and CHO-DCFHP. The average neutralization titers for each group are indicated below each bar. * LOQ, Lower limit of Quantitation.

## Discussion

To date, there remains a large global imbalance in the number of COVID-19 vaccine doses administered, with the high and upper middle income countries administering the vast majority of doses [30]. Various low-cost, more easily manufactured, refrigerator stable, subunit vaccine candidates are being developed to hopefully address this inequity including (1) a receptor binding domain (RBD) antigen formulated with aluminum hydroxide and CpG adjuvants (CORBEVAX™ from Biological E), approved for emergency use in India and Indonesia [31, 32], (2) a spike protein antigen adjuvanted with the saponin-based adjuvant Matrix M (Nuvaxovid™ from Novavax), approved for emergency use by the US FDA [33], (3) a plant-produced VLP of the spike protein antigen adjuvanted with ASO3 (COVIFENZ® from Medicago with adjuvant GSK), approved for use in Canada [34], and (4) a RBD-nanoparticle vaccine antigen designed by IPD adjuvated with ASO3 from GSK (SKYCovine® from SK), which is approved for use in South Korea [35].

Since these recombinant protein antigen-based COVID-19 vaccines employ novel adjuvants, we sought in this work to optimize conventional aluminum-salt adjuvant formulations for the DCFHP nanoparticle to target and facilitate its use in LMICs. Despite the successful development of several new immunologic adjuvants over the past two decades, including the aforementioned COVID-19 vaccines [36], newer adjuvants still must be considered in not only in terms of their safety profile in various patient populations (e.g., adult vs children vs infants), but also their cost, availability of GMP sources, and their compatibility with vaccine antigens during storage within the global vaccine cold chain [24]. To this end, aluminum-salt adjuvanted vaccines have several advantages that make their use attractive such as low cost, wide availability of GMP sources, and decades of safety and efficacy data for their use as adjuvants in healthy infants and children [22, 24, 37]. An additional potential advantage would be the ability to eventually add this new COVID-19 vaccine candidate to pediatric combination vaccines that contain only aluminum adjuvants [38, 39].

### Formulation design to modify extent and strength of antigen-adjuvant interactions and effects on in vivo immunogenicity in mice

Langmuir adsorption isotherm results demonstrated that the DCFHP antigen binds to the aluminum hydroxide (Alhydrogel™, AH) adjuvant with monolayer coverage (adsorptive capacity, Qmax value) consistent with those reported previously with model proteins and other vaccine antigens [40, 41]. The binding strength (adsorptive coefficient, K_L_ value) of DCFHP-AH interaction was in the higher-end range of those observed with other recombinant protein antigens, yet there are examples of other vaccine antigens that bind even more tightly to alum. For example, the HBV vaccine in which the Hepatitis-B surface antigen binds ∼10X more tightly to alum [42].

The molecular mechanisms of DCFHP-AH interaction are mainly electrostatic in nature since pre-treatment of AH with sodium phosphate (which modified the surface charge of colloidal suspension from positive to negative; see results) led to a reduction in the amount of negatively charged DCFHP antigen bound to AH. Additional non-covalent interactions, however, such as hydrophobic interactions likely also play a role, since for example, ∼10% of DCFHP antigen still binds highly negatively charged versions of alum (e.g., Adjuphos, AP or AH treated with 200 mM phosphate). Due to limited material, more quantitative determination of Qmax and K_L_ values of the adsorption of DCFHP to AH in the presence of varying phosphate buffer concentrations was not performed, but is suggested for future work.

Interestingly, similar levels of pseudovirus neutralizing titers were generated in mice independent of the percent of DCFHP antigen bound to AH, yet in the absence of alum adjuvant, much lower immune responses were observed. These results confirm the necessity of alum adjuvant to generate robust neutralizing immune responses, yet the amount of DCFHP bound to the alum adjuvant is not a crucial consideration. Previous reports have highlighted the antigen-specific nature to immunopotentiation after immunization versus percent vaccine antigen bound to aluminum-salt adjuvants. Immunogenicity results have ranged from independent of the amount of antigen adsorbed to alum [43-47] vs. numerous examples of significantly higher antibody titers when vaccine antigens are bound to alum adjuvants [22], e.g., a recombinant Streptococcus pneumoniae vaccine antigen formulated with AH (antigen 100% bound) compared to AP (antigen 0% bound) [48].

### Effect of antigen-adjuvant interactions on the structural integrity and conformational stability of AH-adsorbed DCFHP

DCFHP displayed thermodynamic alterations after adsorption to AH-adjuvant as measured by DSC, e.g., a ∼10°C change in Tm3 values and >2 fold decrease in apparent enthalpy of unfolding (ΔH’) was observed when DCFHP was 100% bound to Alhydrogel compared to solution. We also noted intermediate values for ΔH’ (between the solution and ∼100% AH adsorbed samples) when the extent of the antigen-adjuvant interactions were decreased by the presence of 20-200 mM sodium phosphate. Interestingly, no further effect on Tm3 values were noted with phosphate buffer addition. Several mechanistic studies of model proteins adsorbed to aluminum-salt adjuvants have been reported [40, 49], with structural alterations of these model proteins [40, 49] bound to colloidal suspensions of aluminum-salts could represent either thermodynamically favorable or unfavorable conformations [45], which could manifest as changes in Tm and/or ΔH’ values. In the case of DSC analysis of AH-bound vs solution DCFHP, the reductions in the ΔH’ values are consistent with the increases in the binding strength of DCFHP to AH as modulated by addition of different levels of phosphate buffer.

Previous reports with recombinant protein vaccine antigens interacting with various alum adjuvants demonstrate an antigen specific nature to such binding studies. Agarwal et al. (2020) examined the P[8] antigen of a subunit rotavirus vaccine candidate by DSC and no notable changes in Tm or ΔH’ values were observed after AH adsorption, however, a ∼5°C reduction of the Tonset values was observed [23]. Interestingly, HX-MS analysis of the closely related AH-adsorbed P[4] antigen revealed site-specific structural alterations in the key epitope involved in the binding of a P[4] specific mAb [50]. As another example, Bajoria et al. (2022) performed DSC analysis of a SARS-CoV-2 receptor binding domain (RBD) vaccine candidate before and after adsorption to AH, and results demonstrated a large reduction in both Tm (∼12°C) and ΔH’ values [51]. Finally, when several Neisseria meningitidis recombinant antigens (4CmenB vaccine antigens) were adsorbed to AH-adjuvant and analyzed by DSC (vs solution samples), the AH-adsorbed NadA antigen displayed ∼10°C increase in Tm value, the AH-adsorbed GNA2091 antigen showed a 9°C decrease, and smaller changes (∼ 2-5°C) in Tm values were observed with the other AH-adsorbed antigens [52].

Another commonly observed phenomenon with alum-adsorbed vaccine antigens is that the binding strength (interaction sites) can increase during storage [25] [22]. This was observed for DCFHP-AH interactions since it became more difficult to desorb the antigen using mild conditions as a function of increasing time and temperature. Although varying levels of antigen-adjuvant interactions did not affect mice immunogenicity in this study, further confirmation is warranted to confirm no negative effects on immunogenicity using long-term storage stability samples. In this work, we further explore the nature of the interactions of DCFHP with AH adjuvant by monitoring the ability of the AH-bound DCFHP to bind the ACE2 receptor as described in the next section.

### Effect of antigen-adjuvant interactions of AH-adsorbed DCFHP on ACE2 receptor binding

By developing a competitive ELISA assay to monitor the binding of RBD portion of the DCFHP antigen to the ACE2 receptor, we could evaluate structural alterations of AH-adsorbed DCFHP at the “local key epitope” level. As expected, we observed decreases in ACE2 binding during storage over one month at 25°C, which can be interpreted as conformational alterations at the ACE2 binding site of DCFHP at elevated temperatures. Unexpectedly, however, a small but statistically significant increase in ACE2 binding in the AH-adsorbed DCFHP samples was observed after storage for one month at 4°C.

One possible explanation for this observation is the shift in equilibrium of the spike RBDs from a mixture of “up” and “down” conformations to majority “up” conformation on the surface of the ferritin nanoparticle while adsorbed to Alhydrogel. This would result in more ACE2 receptors that can bind to each spike trimer on the surface the DFCHP antigen. Our analysis from a large dataset of AH-adsorbed DFCHP samples incubated at 4°C for an average of 10 days revealed a statistically significant increase in ACE2 receptor binding when compared to freshly adsorbed samples. These results also suggest that adsorption to AH-adjuvant may promote this shift. Others have observed similar shifts in equilibrium in the up/down conformations in the prefusion stabilized spike protein, albeit not in the presence of an adjuvant [53]. Another, albeit unlikely, possibility for this change is cold temperature induced structural alterations of the spike trimers as observed by Edwards et al (2020) [53]. Further investigation is required to better understand the nature of the structural alterations that occur due to DCFHP adsorption to AH-adjuvant followed by storage at 4°C for a few weeks. To this end, additional competitive ELISA assays with mAbs against other spike protein epitopes is suggested.

Another possible explanation for these observations is a “maturation effect” in which the antigen distribution across the surface of the aluminum-salt changes over time and reaches an equilibrium state over several weeks, as was recently reported by Laera et al. (2023) with AH-binding with several protein antigens in a multivalent vaccine [54]. This maturation effect is not only related to the nature of the antigen and adjuvant, but also the formulation process conditions (e.g., order of addition). The subsequent impact of maturation effects with AH-adsorbed antigens on their interaction with antigen-specific antibodies in a competitive ELISA assay is likely antigen-specific. For example, Sawant et al. (2021) observed increased mAb binding as a function of storage time at 4°C with the P[4] antigen of a subunit rotavirus vaccine candidate adsorbed to AH [55]. In contrast, Sharma et al. (2023) did not observe increased mAb binding of a quadrivalent HPV vaccine adsorbed to AH as a function of storage time at 4°C [56]. In both these examples a similarly designed competitive ELISA format was employed as was used in this work.

### Analytical characterization and comparability assessment of DCFHP produced in two different cell lines

Previous reports describing the design and immunogenicity of the DCFHP antigen [14], as well as the formulation experiments described in this work, were performed using DCFHP antigen produced in Expi293 cells. Although useful for rapidly producing small amounts of material in the laboratory for initial experiments, the commercial production of recombinant viral glycoproteins in stable cell lines such as CHO cells is the industry gold standard due to increased yields, reproducibility, and ease of scalability [57-59]. To better translate the findings of this work to clinical use, we performed an analytical comparability study, combined with mouse immunogenicity studies, to assess the quality of the DCFHP antigen produced in Expi293 and CHO cells. Manufacturing process changes, especially cell line changes, often lead to alterations in macromolecule structure and biological activity that require careful evaluation to ensure a comparable product is produced [60, 61].

To this end, a second major goal of this work was to characterize and compare DCFHP that was produced in Expi293 and CHO cells. As part of this analyses, we also identified the most promising physiochemical methods for determining the critical quality attributes of the DCFHP antigen. DCFHP produced in either cell lines were of the same primary structure and fully assembled into nanoparticles at the expected molecular size with minimal aggregates, maintained ACE2 binding ability, and induced high pseudovirus neutralizing titers in mice. We did, however, observe differences in the N-linked glycosylation patterns, which was not unexpected and has been previously observed by others [26]. These glycosylation differences did not affect physicochemical properties, *in vitro* potency (binding the ACE2 receptor) or *in vivo* immunogenicity (in mice). Recombinant glycoprotein production using CHO cells is the industry gold standard because of high yields (in g/L range), stable gene expression, and relative ease of scalability [58, 59].

### Conclusions and Future Work

In this work, a combination of analytical characterization and formulation development studies were performed to address an often-overlooked area of subunit vaccine development: the effect of adjuvants not only the immunogenicity, but also the pharmaceutical properties and storage stability of the formulated antigen. To this end, a major focus was to better understand the inter-relationships between antigen-adjuvant interactions, pharmaceutical properties, and various potency measurements. We demonstrate DCFHP is poorly immunogenic in mice without an adjuvant and immune responses are greatly boosted using conventional aluminum-salt adjuvant (Alhydrogel, AH). Interestingly, the percent of antigen bound to the AH adjuvant did not affect immunogenicity as measured the pseudovirus neutralization titers. These results indicate that loose association and/or co-administration of DCFHP with the AH is sufficient to drive a robust immune response.

In terms of pharmaceutical stability profiles of AH-adjuvanted DCFHP, we focused this work on short-term stability assessments over one month of storage. During storage at 4°C, a “maturation effect” in the interactions between the DCFHP antigen and the AH adjuvant was observed leading to enhanced ACE2 receptor binding with concomitant decreased ability to desorb the antigen from AH adjuvant. In contrast, at elevated temperatures (25°C), greater structural rearrangements of the DCFHP antigen occur leading to notable losses of ACE2 receptor binding and even greater difficulty in desorbing the antigen from the AH. Finally, under more extreme thermal stress conditions encountered by temperature ramping DSC experiments, DCFHP-Alhydrogel interactions affect the overall conformational stability of the DCFHP antigen as observed by decreases in one of the thermal melting temperatures (Tm3) values as well as the overall apparent enthalpy values (compared to DCFHP in solution). Future studies will focus on better understanding of the long-term stability profiles of AH-adsorbed DCFHP vaccine candidate during refrigerator and room temperature storage. To this end, the physicochemical methods and potency assays described in this work can be employed. Ambient temperature stability could be a major advantage in easing worldwide vaccine distribution and assist in overcoming vaccine inequity.

Finally, we developed an overall analytical comparability assessment strategy, combining physiochemical methods with potency assays including *in vitro* ACE2 receptor binding and *in vivo* mouse immunogenicity. Results not only demonstrated comparability of DCFHP antigen produced in Expi293 and CHO cells, but also identified analytical assays for future use to monitor lot-to-lot consistency and ensure the quality of this SARS-CoV-2 spike ferritin nanoparticle vaccine candidate during future process development and scaleup.

## Abbreviations

AH: Alhydrogel™
AP: Adjuphos™
BLI: Bio-Layer interferometry
CD: Circular dichroism
CMC: Chemistry, Manufacturing, and Control
DCFHP: Delta-C70-Ferritin-HexaPro
DLS: Dynamic light scattering
dPBS: Dulbecco’s phosphate-buffered saline
DSC: Differential scanning calorimetry
LDS: Lithium dodecyl sulfate
SARS-CoV-2: Severe acute respiratory syndrome coronavirus 2
SEC-MALS: Size exclusion chromatography with multi-angle light scattering detection
SV-AUC: Sedimentation velocity analytical ultracentrifugation
NHP: Non-human primates

## Acknowledgements-

We thank J. Bloom and A. Greaney for plasmids and cells related to viral neutralization assays. This work was supported by the Frank Quattrone & Denise Foderaro Family Research Fund, the Chan Zuckerberg Biohub, the Stanford Innovative Medicines Accelerator, the Virginia & D.K. Ludwig Fund for Cancer Research, and an NIH Director’s Pioneer Award (DP1AI158125) to P.S.K.

