## Supplemental Material for "Formulation development and comparability studies with an aluminum-salt adjuvanted SARS-CoV-2 Spike ferritin nanoparticle vaccine antigen produced from two different cell lines"

### Supplemental Materials and Methods

**Materials**- Spike-Ferritin nanoparticles (DCFHP) were an updated version of SΔC-Fer, previously described by Powell et al., [1], containing four additional proline amino acid substitutions F817P, A892P, A899P, A942P and a mutated S1/S2 furin cleavage site as described in [2]. The nanoparticles were produced in Expi293 cells as described [1], with minor modifications. Briefly, the gene encoding DCFHP was expressed in Expi293F cells cultured in media containing Freestyle and Expi media (ThermoFisher) mixed at 2:1 ratio and transfected with FectPro reagent (VWR). After 4-5 days of culture, the nanoparticles' containing media was clarified by centrifugation and filtration. Clarified media was diluted two-fold with 20 mM Tris pH 8.0 followed by addition of NaCl to 200nM final concentration. The sample was loaded on the HiTrap Q (Cytiva) column pre-equilibrated with 20 mM Tris, 200 mM NaCl, pH 8.0 using a sample pump and the unbound fraction that contained the nanoparticles was collected. The sample was washed twice by concentrating with centrifugal concentrators (Amicon, 100 kDa MWCO), diluting with 20 mM Tris, 150 mM NaCl, pH 8.0 and concentrated again. Two mL of sample was injected on a AKTA pure FPLC (GE Healthcare) using a 2mL loop and run over the SRT1000 gel filtration column, which was pre-equilibrated 20 mM, 150mM, Tris pH 7.5.

DCFHP produced in CHO cells was filtered with 0.22 µm filter, dialyzed overnight against 20 mM Tris, 150mM NaCl, pH 7.5 using 1 MDa dialysis unit (Spectra/por Float-A-Lyzer G2 Dialysis), filtered again and further purified on AKTA pure FPLC with a SRT1000 gel filtration column pre-equilibrated in 20 mM Tris, 150mM NaCl, pH 7.5. All chemicals and excipients were purchased from Sigma (St. Louis, MO) except for sucrose (purchased from Pfanstiehl Inc.,

Waukegan, IL) and Alhydrogel (purchased from InvivoGen, San Diego, CA.) All chemicals were of high purity (≥99%).

**Size-Exclusion Chromatography-MALS (SEC-MALS)**- A Shimadzu Prominence UFLC HPLC system equipped with a diode array detector, a Wyatt Optilab-rEX refractive index detector, and a Wyatt DAWN HELEOS II multi-angle light scattering detector was used. DCFHP samples were diluted to 0.2 mg/mL in 20mM Tris, 150mM NaCl, 5% Sucrose, pH 7.5 and 25µl and were injected onto a Sepax, SRT SEC-1000, 5 µm 1000A, 4.6x300 mm size exclusion column. The column was operated at 30°C and equilibrated with at least 10 column volumes of mobile phase (0.2 M sodium phosphate, pH 6.8) prior to sample injection. A flow rate of 0.35 ml/min was used with a 30 min run time. LC solutions software (Shimadzu) for was used for data analysis. The MALS data was analyzed using Astra v6.1 (Wyatt technologies) fitting all data to the Zimm model using BSA as a normalization standard.

**Far-UV Circular Dichroism-** Circular Dichroism was performed using a Chirascan-Plus ACD spectrometer (Applied Photophysics Ltd, Leatherhead UK) equipped with an autosampler fitted with a 1mm temperature-controlled quartz flow cell. DCFHP samples were diluted to 0.1 mg/mL in 20mM Tris, 150mM NaCl, 5% sucrose, pH 7.5. The light source, monochromator, and flow cell sample holder were continuously purged with $N_2$. Samples were equilibrated at 10°C and measured from 260-200 nm using 1 nm step and a 1 s integration time. All spectra were subjected to a 5 point Savitzky-Golay smoothing filter. Ellipticity of the buffer alone was subtracted from all sample measurements.

**Intrinsic Fluorescence spectroscopy**- The intrinsic tryptophan fluorescence of **DCFHP** samples were measured using a dual emission PTI QM-40 Spectrofluorometer (Horiba, Birmingham, NJ) equipped with a 4-position cell holder peltier temperature control device, a high power continuous 75 W short-arc Xe lamp (Ushio), and a Hamamatsu R1527 photomultiplier tube. Data were collected using FelixGX software. Fluorescence emission spectra of 0.1 mg/mL DCFHP were recorded using an excitation wavelength of 295 nm (>95% Trp) and monitoring emission from 310-390 nm with a step size of 1 nm and an integration time of 1 s.

**Differential Scanning Calorimetry**- DCFHP samples were diluted to 0.2 mg/mL in 20mM Tris, 100mM NaCl, 5% sucrose, pH 7.5 and loaded in a DSC autosampler tray held at 4°C. DSC was performed using an Auto-VP capillary differential scanning calorimeter (Malvern, Northhampton, MA). Sample and reference cells pressurized with $N_2$ at ~65 psi. Two water-water scans were taken prior to the first reference scan. Samples were heated from 10-100°C using a scan rate of 60°C/hr, a pre-scan thermostat of 15 min, no feedback gain, and a filtering period of 10 points/s. Reference subtraction, baseline correction, and concentration normalization were performed using the instrument software. Since the baseline noise was significantly increased in the Alhydrogel containing scans, these scans were subjected to a 38-point Savitzky-Golay smoothing filter.

**Sedimentation Velocity Analytical Ultracentrifugation**- Sedimentation velocity (SV-AUC) experiments were performed using an Optima analytical ultracentrifuge equipped with a scanning ultraviolet-visible optical system (Beckman Coulter, Indianapolis, IN.) A rotor speed of 7,000 rpm, rotor temperature of 20°C, UV detection at 280 nm, and a scan frequency of 60 seconds were used. DCFHP samples (at 0.2 mg/ml) in formulation buffer (20mM Tris, 150mM NaCl, 5% Sucrose, pH 7.5) alone were loaded into Beckman charcoal-epon two sector cells with 12 mm centerpieces and either sapphire or quartz windows and ultracentrifugation was performed for 800 total scans. Sedimentation data were analyzed using Sedfit (Peter Schuck, NIH) using a continuous c(s) model in the range of 0 to 100 svedbergs. A resolution of 300 points per distribution and a confidence level of 0.95 were applied to all fits. Baseline, radial independent noise, and time independent noise were fit, while the meniscus and bottom positions were set manually. The c(s) distributions were imported into Origin 2018 (OriginLab, Northampton, MA) for peak integrations and graph generation.

**Dynamic Light scattering**- Dynamic light scattering was performed using a Wyatt DynaPro 3 plate reader. Twenty-five microliters were loaded in each well of a 384 well plate (Corning) at a final protein concentration of 0.1 mg/mL DCFHP. After brief centrifugation, DLS was performed at 25°C with automatic laser attenuation enabled. All DLS data was corrected for solution viscosity and temperature.

**Peptide Mapping Analysis**- Prior to LCMS analysis, N-glycans from DCFHP were removed following an overnight 37°C incubation with EndoH or PNGaseF (New England Biolabs). DCFHP peptides were then generated using a commercial kit (S-Trap$^{TM}$ micro kit, Protifi LLC). Briefly, 20 µg of DCFHP was reduced with TCEP, heat denatured (10 min at 98°C), and then alkylated with methyl methanethiolsulfonate (MMTS). DCFHP was then treated for 2 hrs at 47°C with trypsin (Promega Corporation) or Chymotrypsin (ThermoScientific) and injected onto an Advanced Peptide column (2.1 x 150 mm, 2.7 µm, Agilent Technologies). The LC gradient of consisted of 2-45% B (A: 0.1% formic acid in water, B: 0.1% formic acid in acetonitrile) over 73 min at a flow rate of 0.3 mL/min. Eluting peptides were then mass analyzed using a 6545XT QTOF (Agilent Technologies). Mass spectra were collected from 250-1700 m/z at 1 spectra/sec. The threshold for MS/MS analysis was 50,000 counts and the two most abundant ions were selected for CID fragmentation per cycle. Mass spectra were processed using MassHunter Bioconfirm v10.0 software (Agilent Technologies).

**N-linked Glycan Oligosaccharide Mapping Analysis-** Approximately 15 µg of the glycoprotein was de-glycosylated and the released N-linked glycans were labeled and extracted using a commercial kit (GlycoWorks *Rapi*Fluor-MS *N*-Glycan Kit, Waters Corporation). One µl of the *Rapi*Fluor-labeled glycans was injected into 1290 Infinity II UHPLC system (Agilent Technologies) containing a 2.1 x 150 mm, 1.8 µm Glycan column (Agilent Technologies) maintained at 40°C. The LC mobile phases consisted of water with 50 mM ammonium formate (mobile phase A) and acetonitrile (mobile phase B). Glycan separation was achieved using a 36 min 75-63% mobile phase B gradient. Elution of the glycans was monitored using either an in-line fluorescence detector ($\lambda_{EX}$: 265 nm; $\lambda_{EM}$: 425 nm) or an in-line 6545XT QTOF mass spectrometer (Agilent Technologies). The electrospray ionization parameters consisted of: 150°C gas temperature, 3000V Vcap, and 100V fragmentor. Mass spectra were collected from 600-1700 m/z at 1 spectra/sec. Mass spectra were processed using MassHunter Bioconfirm v10.0 software (Agilent Technologies).

**Bio-Layer interferometry-** Binding of DCFHP with ACE2-Fc was determined using an Octet Red96 Biolayer Interferometry System (Pall Forte Bio LLC, Fremont, CA). Binding experiments were performed in triplicate using Protein G biosensors (Forte Bio, Cat No. 18-5082) in 96-well black microplates (Greiner Bio-One). Assay kinetics buffer (Quality Biological, 1X PBS pH 7.2 + 0.05% PS-80) was used for all baseline, dissociation, and reference wells as well as for diluting the ACE2-Fc receptor to loading concentration of 10 mcg/mL, and DCFHP at a starting concentration of 25 mcg/mL, followed by a 7-point 1:2 serial dilution in the kinetics buffer. Biosensors were hydrated for ~10 min in kinetics buffer prior to the run. A typical run comprised of a baseline step (60 sec), followed by loading step (300 sec), another baseline step (60 sec), association step (250 sec), and lastly, dissociation step (1800 sec) with shake speed maintained at

1000 rpm throughout the experiment. Data analysis was performed using Octet Data Analysis software (v 10.0, Forte Bio). Following reference subtraction, baseline alignment, inter-step correction, and data processing with Savitzky-Golay filtering. Since there was negligible dissociation, Kd calculations were not possible.

**Competitive ACE2 ELISA**- DCFHP samples were diluted to 25-50 mcg/mL in 20mM Tris, 150mM NaCl, 5% Sucrose, pH 7.5 (in solution and AH-adsorbed) and were incubated in casein blocking buffer (Thermo-Fisher) with 0.05% Tween-20 such that the blocking buffer made up 1/3 of the final volume for one hour at room temperature with end over end rotation. Samples were then transferred to 96-well PCR plates (Thermo-Fisher) and 1:2 serial dilutions were made across the plates followed by incubation with ACE2-Fc receptor [3] (Institute for Protein Design, Seattle, Washington) at 0.04 mcg/ml. Plates were sealed using strip caps and incubated overnight at ambient temperature with gentle end over end rotation. Plates were then centrifuged at 1,600 x for 5 min, and the supernatant containing unbound ACE2-Fc was transferred to 96-well polysorp Nunc Immobilizer Amine or Maxisorp assay plates (Thermo-Fisher) coated with recombinant SARS-CoV-2 receptor binding domain (RBD) [4] (Institute for Protein Design, Seattle, Washington) at 1 mcg/ml in dPBS. Prior to this step, the assay plate was blocked for 1 hour at 25°C in the same casein blocking buffer with 0.05% Tween-20. The plate was then incubated for 2 hours at 25°C and the amount of bound ACE2-Fc on the plate was detected using a 1:5000 dilution of anti-human Ig HRP–conjugated secondary antibody (Southern Biotech). Plates were developed for color formation using Tetramethylbenzidine substrate solution (Sigma-Aldrich, St. Louis, MO) by incubating at room temperature in dark for 12 min and quenched using 1N N $H_2SO_4$. Optical density (OD) at 450 nm was measured using SpectraMax iD3 (Molecular Devices, San Jose, CA) and the data were analyzed using Origin software (OriginLab, Northampton, MA) after blank subtraction. The concentration was calculated from the OD 450 values using parameters obtained from a 4-point logistic fit from the standard run on each plate with no weighting of individual data points. Between each step (until the development and quench step) the assay plates were washed three times with 0.3 mL dPBS w/ 0.05% Tween-20 using a Bio-Tek automated plate washer (Agilent Technologies, Santa Clara, CA.)

**Preparation of AH-adjuvanted DCFHP samples for Antigen-adjuvant binding, *in vitro* stability, and *in vivo* mouse studies-** Binding of DCFHP to Alhydrogel® (AH) was performed by dilution of DCFHP in 20mM Tris, 150mM NaCl, 5% sucrose, pH 7.5 with a final Aluminum concentration of 1.5 mg/mL (Invitrogen, San Diego, CA). Samples were incubated for 1 hour at room temperature with gentle end over end rotation, and adsorption to the aluminum salt was determined by centrifugation of samples at 4,000 x g for 5 min and analysis of the supernatant by either UV-Visible spectroscopy or SDS-PAGE. Adjuvanted DCFHP containing formulations were filled in pre-sterilized 2 mL glass vials (Fiolax clear type, Schott), capped with pre-sterilized FluroTec® coated 13 mm serum stoppers, and sealed using 13mm aluminum seals (West Pharmaceuticals). All vials were filled under aseptic conditions in a Class II Biosafety cabinet (Labconco). For the mouse immunogenicity studies, the vials were shipped overnight on wet ice from Lawrence, KS to Palo Alto, CA. Stability samples were stored in temperature-controlled stability chambers (Torrey-Pines scientific).

The desorption of the DCFHP antigen from the AH adjuvant was performed either by using mild or strong conditions: (1) Mild desorption was performed by pelleting the AH-bound DCFHP as described above and resuspending the pellet in 0.4 M sodium phosphate, pH 7.5 followed by incubation at 37°C for 30 minutes. The solution was then centrifuged to pellet the AH and the supernatant was analyzed by SDS-PAGE or UV-Visible spectroscopy, (2) Strong desorption was performed by pelleting the AH-bound DCFHP as described above and resuspending the pellet in strong desorption buffer containing 0.4 M sodium phosphate, 50mM DTT (Bio-Rad) and 1X LDS loading buffer at pH 7.6 (10% glycerol, 1% lithium dodecyl sulfate (LDS), 0.2 M triethanolamine-Cl, 1% Ficoll®-400, 0.0125% phenol red, 0.0125% Coomassie G250, 0.5 mM EDTA disodium; Thermo-Fisher), and heating at 95°C for 5-10 min. The solution was then centrifuged to pellet the aluminum salt, followed by analysis of the supernatant by SDS-PAGE. Estimation of the protein concentration in the sample supernatant by SDS-PAGE was determined by performing densitometry on known amounts of DCFHP using ImageJ software (NIH, Bethesda, MD, available at: https://imagej.nih.gov/ij/). A standard curve was constructed using 1.5-0.25 mcg DCFHP standards by graphing the densitometry values as a function of known mass and performing linear regression using Origin 2018 software (OriginLab, Northampton, MA).

**Langmuir adsorption isotherms**- Binding isotherms were constructed to calculate the adsorptive capacity (Qmax) and binding strength ($K_L$) values of AH-bound DCFHP. Varying amounts of DCFHP (0-600 µg/mL) was incubated with a fixed amount of AH (50 µg Aluminum) using 20mM Tris, 150mM NaCl, 5% sucrose, pH 7.5 as the diluent. The mixtures were incubated at 25°C for one hour with gentle end over end rotation, samples were then centrifuged at 4,000 x g for 5 minutes, and the supernatant was measured by UV-Visible spectroscopy using a Lunatic UV-Vis instrument (Unchained Labs) with the protein concentration calculated using the extinction coefficient (1.014 $cm^{-1}$ $M^{-1}$) calculated from the primary amino acid sequence. The protein concentration of the supernatant (Ce) was calculated using the A280 value after light scattering correction of the raw spectra. The data were then analyzed to fit to the linearized form of the Langmuir equation to calculate the Qmax and $K_L$.

The Langmuir equation is shown below

$$q_e = \frac{Q_{max} K_L C_e}{1 + KL\, C_e}$$

The linearized form of the Langmuir equation:

$$\frac{C_e}{q_e} = \frac{1}{Q_{max}} C_e + \frac{1}{K_L Q_{max}}$$

Where $q_e$ and $C_e$ are solid phase and liquid phase equilibrium concentrations, respectively. $Q_{max}$ is maximum (monolayer) binding capacity. $K_L$ represents the Langmuir isotherm constant, which represents binding strength [5].

**Mice Immunization studies-** Balb/C mice were procured from Jackson Laboratories (Bar Harbor, ME). All animals were maintained at Stanford University according to Public Health Service Policy for 'Humane Care and Use of Laboratory Animals' following a protocol approved by Stanford University Administrative Panel on Laboratory Animal Care (APLAC). Eight to ten weeks old female Balb/C mice were immunized by intramuscular injection of 10 mcg of DCFHP with different formulations of aluminum salt adjuvant (Alhydrogel, InvivoGen, San Diego, CA) (Supplemental Table S3)**.** Blood samples were collected from the immunized mice in a micro tube containing Z-Gel (Sarstedt AG and Company) via bleeding of retro-orbital plexus. Serum sample was isolated by centrifugation of the collection tube at 10,000 x *g* for 5 min and stored at -80$^{o}$C. Viral neutralizing titers of the serum samples were determined by the method described below.

**SARS-CoV-2 pseudo-typed lentivirus production and viral neutralization assays-** SARS-CoV-2 spike pseudo-typed lentivirus was produced in HEK293T cells using BioT reagent (Bioland Scientific LLC). Five million cells were seeded in D10 medium (DMEM + additives: 10% fetal bovine serum, L-glutamate, penicillin, streptomycin, and 10 mM HEPES) in 10-cm plates one day prior to transfection. A five-plasmid system [6] was used for viral production: the lentiviral packaging vector (pHAGE_Luc2_IRES_ZsGreen), the SARS-CoV-2 spike, and lentiviral helper plasmids (HDM-Hgpm2, HDM-Tat1b, and pRC-CMV_Rev1b). The spike vector contained the C-terminal deletion of 21 amino acids (D21) of wild-type spike sequence from the Wuhan-Hu-1 strain of SARS-CoV-2 (GenBank NC_045512) or from SARS-CoV-2 B.1.1.529.1 (Omicron) variant. Plasmids were added to D10 media as follows: 10 µg pHAGE_Luc2_IRS_ZsGreen, 3.4 µg SARS-CoV-2 spike, 2.2 µg HDM-Hgpm2, 2.2 µg HDM-Tat1b, and 2.2 µg pRC-CMV_Rev1b in a final volume of 1 mL. To form transfection complexes, 30 µL of BioT reagent were added (Bioland Scientific LLC). Transfection reactions were incubated for 10 min at room temperature, and the volume was made up to 10 mL. The media was removed from the seeding plate, followed by dropwise addition of transfection mixtures to the plated cells. Medium was removed ~24 h post transfection and replaced with 10 mL of fresh D10 medium. Virus-containing culture supernatants were harvested ~72 h post transfection via centrifugation at 300 x *g* for 5 min and filtered through a 0.45-µm filter. Finally, the supernatant was buffered with final concentration of 10 mM HEPES. Viral stocks were aliquoted and stored at -80 °C until use.

For viral neutralization assays, TMPRSS2/ACE2/HeLa [3] cells were plated in white-walled clear-bottom 96-well plates at 10,000 cells/well one day prior to infection. Mouse serum samples were heat inactivated for 30 min at 56 °C and diluted in D10 medium. Virus was diluted in D10 medium, supplemented with polybrene at a final concentration of 5 µg/mL, and then added to the diluted serum. After incubation, medium was removed from cells and replaced with an equivalent volume of serum and virus mixture and incubated at 37 °C for ~48 hrs. Cells were lysed by adding BriteLite (Perkin Elmer) assay solution and luminescence values were measured with a TECAN Infinite M Plex plate reader. Serum from each animal at each time point was assessed in duplicate with a 6-point serial dilution (starting at 1:100 with 10-fold dilutions). Each assay was performed again in a separate experimental replicate. Each plate was normalized by averaging RLUs from wells with cells only (0% infectivity) and virus only (100% infectivity). Normalized values were fit with a three-parameter non-linear regression inhibitor curve in GraphPad Prism 8.4.1 to obtain $NT_{50}$ (50 % Neutralizing Titer) values. Fits for all serum neutralization assays were constrained to have a value of 0% at the bottom of the fit. The limit of quantitation for this assay is approximately 1:100 serum dilution. Serum samples that failed to neutralize or that neutralized at levels higher than 1:100 was set at the limit of quantitation for statistical analyses. Plots are shown starting at the reciprocal serum dilution of the limit of quantitation ($10^2$).

**Supplemental Table S1. N-glycan analysis of DCFHP produced in Expi293 cells.** The ten most abundant N-glycans are displayed.

| Glycan | Average Relative Fluorescence Peak Area Abundance ± 1SD (%) | Average Relative Ion Abundance ± 1SD (%) | Structure |
|---|---|---|---|
| H3N4F1 | 21.4 ± 0.5 | 24.5 ± 0.1 | 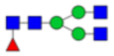 |
| H5N2 | 13.1 ± 0.0 | 12.6 ± 0.1 | 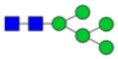 |
| H3N5F1 | 9.6 ± 0.3 | 10.6 ± 0.0 | 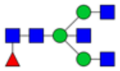 |
| H4N4F1 | 6.4 ± 0.1 | 7.2 ± 0.1 | 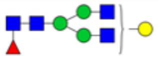 |
| H3N3F1 | 5.6 ± 0.1 | 5.6 ± 0.0 | 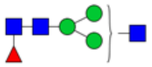 |
| H4N4F1S1 | 10.2 ± 0.1* (elutes at same time as H5N4F1) | 5.4 ± 0.0 | 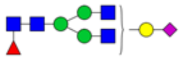 |
| H4N4F1 | 6.4 ± 0.1 | 3.5 ± 0.0 | 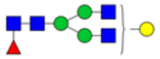 |
| H4N5F1 | 2.0 ± 0.1 | 2.8 ± 0.0 | 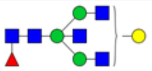 |
| H5N4F1 | 10.2 ± 0.1* (elutes at same time as H4N4F1S1) | 2.8 ± 0.0 | 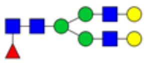 |
| H7N2 | 4.4 ± 0.1 | 2.6 ± 0.0 | 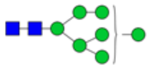 |

**Supplemental Table S2. N-glycan analysis of DCFHP produced in CHO cells.** The ten most abundant N-glycans are displayed.

| Glycan | Average Relative Fluorescence Peak Area Abundance ± 1SD (%) | Average Relative Ion Abundance ± 1SD (%) | Structure |
|---|---|---|---|
| H5N4F1 | 13.8 ± 0.3 | 14.1 ± 0.1 | 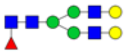 |
| H5N2 | 10.7 ± 0.4 | 13.8 ± 0.2 | 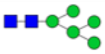 |
| H5N4F1S1 | 10.9 ± 0.2 | 10.9 ± 0.2 | 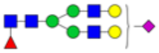 |
| H4N4F1 | 4.4 ± 0.0 | 4.2 ± 0.0 | 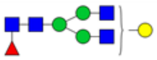 |
| H6N2 | 3.0 ± 0.1 | 4.2 ± 0.0 | 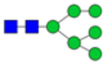 |
| H6N5F1 | 5.1 ± 0.1 | 4.2 ± 0.1 | 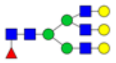 |
| H7N2 | 3.6 ± 0.1 | 4.1 ± 0.0 | 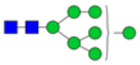 |
| H5N4 | 3.4 ± 0.0 | 3.8 ± 0.1 | 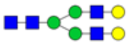 |
| H6N5F1S1 | 4.8 ± 0.1 | 3.8 ± 0.1 | 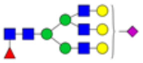 |
| H5N4F1S2 | 4.5 ± 0.1 | 3.3 ± 0.1 | 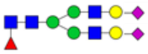 |

### Supplemental Table S3- Summary of samples used in mouse immunogenicity studies

| Formulation | Final DCHFP concentration (mcg/mL) | Antigen amount (per 0.1 mL) (mcg) | Aluminum concentration (mg/mL) | Aluminum amount (per 0.1 mL) (mcg) | IM route injection volume (mL) | % Antigen bound to AH |
|---|---|---|---|---|---|---|
| AH | 100 | 10 | 1.5 | 150 | 0.1 | 100% |
| AH+20mM sodium phosphate | 100 | 10 | 1.5 | 150 | 0.1 | ~40% |
| AH+200mM sodium phosphate | 100 | 10 | 1.5 | 150 | 0.1 | ~10% |
| No adjuvant | 100 | 10 | 0 | 0 | 0.1 | N/A |

**Supplemental Figure S1. Peptide sequence coverage map of Expi293 and CHO produced DCFHP.**

| | CHO | Expi293 |
|---|---|---|
| Overall Sequence Coverage (not including signal peptide) | 94% | 92% |
| Sequence Coverage of Spike Portion | 93% | 91% |
| Sequence Coverage of Ferritin Portion | 98% | 98% |

**CHO DCFHP**
**Expi293 DCFHP**
*Ferritin*

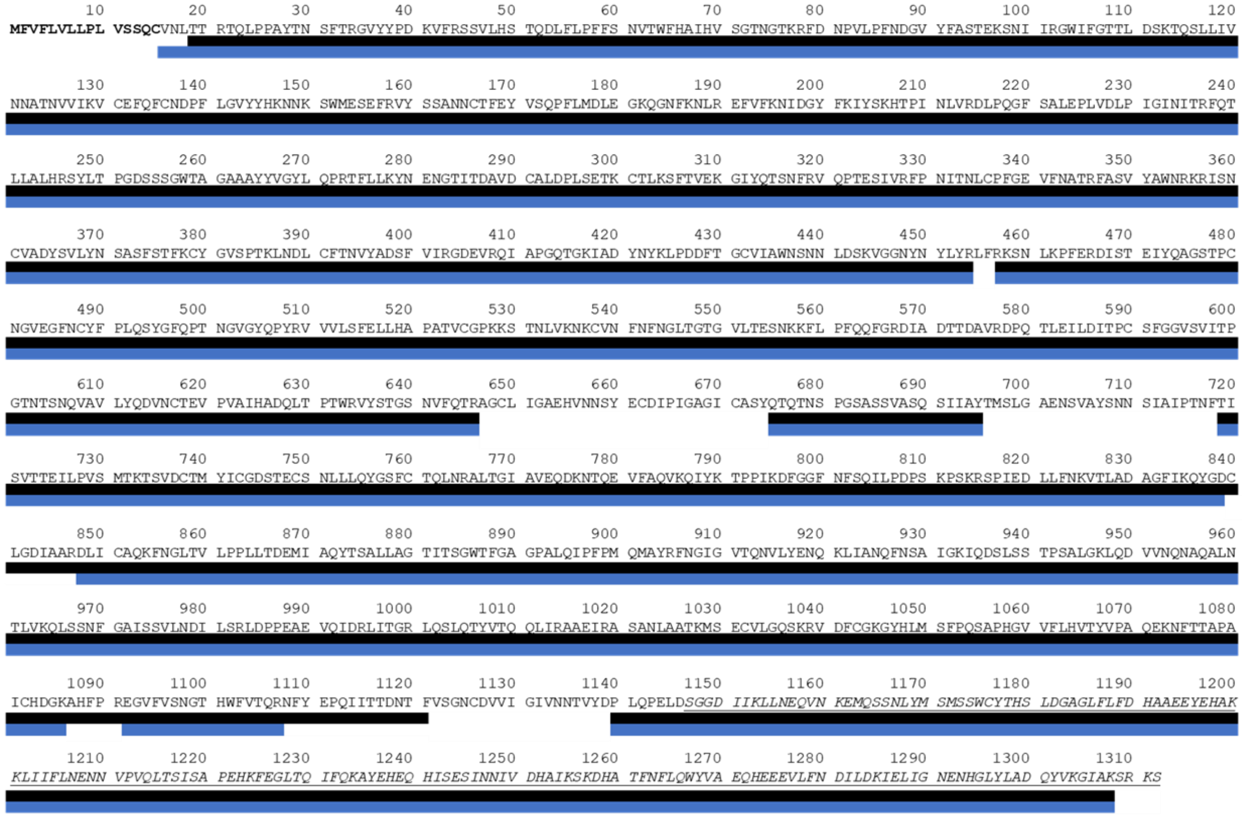
